# Sexual coercion in a natural mandrill population

**DOI:** 10.1101/2022.02.07.479393

**Authors:** Nikolaos Smit, Alice Baniel, Berta Roura-Torres, Paul Amblard-Rambert, Marie J. E. Charpentier, Elise Huchard

## Abstract

Increasing evidence indicates that sexual coercion is widespread. While some coercive strategies are conspicuous, such as forced copulation or sexual harassment, less is known about the ecology and evolution of intimidation, where repeated male aggression promotes future rather than immediate mating success with targeted females. Although known in humans, intimidation was recently reported in chimpanzees (*Pan troglodytes*) and chacma baboons (*Papio ursinus*), where males are regularly violent against females. Here, we investigate the nature of male coercive strategies in wild mandrills (*Mandrillus sphinx*), a primate living in large polygynandrous groups where severe male aggression towards females is rare and females can form coalitions against males. Yet, we found support for all three predictions of the sexual coercion hypothesis, namely that male aggression (1) specifically targets sexually receptive females, (2) inflicts costs to these females, and (3) increases male mating success in the long-term. These results hold true when considering only non-physical threats, or only severe aggression. Finally, we show that high-ranking females are most targeted by males, probably because of their higher reproductive performances, while high-ranking males are most coercive. These results indicate that sexual intimidation is widespread in sexually dimorphic and group-living mammals, and that males and females vary in their propensities to use, and to be exposed to sexual coercion, respectively.

## 1 Introduction

The diverging evolutionary interests of males and females often lead to sexual conflict. While female reproductive success is typically limited by the elevated costs of reproduction, e.g. gestation and lactation in mammals, male reproductive success is primarily determined by the number of mating partners [1]. In some species, males use sexual coercion towards females, defined as “the use by a male of force, or threat of force, that functions to increase the chances that a female will mate with him at a time when she is likely to be fertile, and to decrease the chances that she will mate with other males, at some cost to the female” [2], to improve their mating success [2, 3].

Behavioural ecologists have traditionally documented coercive strategies that are immediately visible, such as forced copulation (when a female is physically restrained by a male to mate with him), sexual harassment (when aggression immediately precedes copulation and is directed until the female cedes; [2]) and coercive mate-guarding (when a male aggressively herds females and enforce close proximity to prevent them to copulate with rival males; [4, 5]). These forms of sexual coercion have been reported from insects [6, 7] to vertebrates [8, 9, 10, 11, 12]. In contrast, longterm forms of sexual coercion – when aggression does not translate immediately but subsequently into mating benefits for the aggressor – are more elusive and have been less studied outside of human societies. Sexual intimidation, when repeated male aggression aims at enforcing future female sexual compliance, has only been documented in two primate societies characterized by severe male aggression to females (chimpanzees (*Pan troglodytes*): [13]; chacma baboons (*Papio ursinus*): [14]). Similarly, males of different taxa (e.g. birds and primates including humans) can also punish females following copulations with rival males to prevent cuckoldry in the future [15, 16, 17, 18].

Sexual coercion is increasingly recognized as a driving force influencing the evolution of mating and social systems in animals [19, 2, 20], including humans [21, 22]. In mammals, male coercive tactics appear most common in polygynous and polygynandrous species where males compete intensively over mating opportunities and a substantial fraction of males fails to secure copulations, and where sexual size dimorphism is pronounced, allowing males to threaten or harass females at low costs [23, 24]. In these species, female impediment to male copulation attempts has been associated with an increased risk of severe injury or even death [25]. The forms of coercion used by males are then likely to vary according to the stability of male-female associations and male dominance status. Short-term strategies such as sexual harassment and forced copulations may be frequently used in solitary species, where males and females only encounter each other for mating [3]. By contrast, long-term strategies, such as intimidation and punishment, are more likely to evolve in species living in stable bisexual groups where males and females maintain medium-to long-term social relationships. Furthermore, in polygynous groups, harassment and forced copulations might be used more frequently by subordinate males that are excluded from mating opportunities [26, 27] while long-term male coercive strategies might be used more often by dominant males to constrain female promiscuity and impose closer proximity (e.g. [28]).

Primates are good candidates to study sexual coercion because the diversity of their social and mating systems may promote various male and female sexual strategies, while their extensive cognitive abilities, including individual recognition and long-term memory, may facilitate the use of long-term male coercive strategies [22]. Such strategies are also promoted by the fact that many primates live in stable bisexual groups where males and females maintain differentiated relationships, and by a widespread male-biased sexual dimorphism associated with polygynous or some polygynandrous mating systems.

In this study, we examine whether males exert sexual coercion in a large natural, polygynandrous group of mandrills (Mandrillus sphinx), a primate from the Cercopithecidae family characterized by an extreme sexual dimorphism in body size (males are 3.4 times heavier than females; [29]) and canine length [30]. Mandrills are seasonal breeders and most males immigrate in the social group at the onset of the mating season ([31]; which generally lasts every year from April to September [32]), resulting in intense male-male mating competition [33]. Male reproductive skew is high, since the alpha male monopolizes 60-70% of reproductions [34, 35]. Female mandrills develop perineal swellings during fertility that grow in size as they approach ovulation and dominant males focus their mate-guarding efforts on maximally swollen females [36]. Yet, both sexes mate promiscuously and females may exhibit some forms of mate choice [37], for example by avoiding males’ attempts to copulate or interrupting copulation before ejaculation (MJEC personal observation). Severe male aggression towards females occurs but appears relatively infrequent for human observers. Female relatives form tight social relationships [34], including aggressive coalitions against males that can, exceptionally, lead to male’s death (in captivity: [38]). Studying male sexual coercion in this species, where most males are temporary residents in the group during the mating season, females can retaliate against males and severe male aggression against females is inconspicuous, appears thus highly relevant.

We test the three key predictions of the sexual coercion hypothesis [2], namely that male aggression (i) targets sexually receptive females more than females in other reproductive states, (ii) is costly to females in the form of a greater exposure to injuries, and (iii) increases male mating success with the victim. For this last prediction, we further investigate different forms of coercion by testing if aggression by a male towards a female increases his chances to mate with her within the following minutes (harassment) or within a longer time-window (intimidation). We also test whether a female that has just copulated with a given male receives immediate aggression from other male(s) as a punishment. We subsequently test whether higher-ranking males are more aggressive towards females during the mating season given the high reproductive skew in their favour. Finally, as an alternative hypothesis to sexual coercion, we test the “aggressive male phenotype” hypothesis, stating that the correlation between male aggression and mating is observed because females prefer to copulate with aggressive males due to direct (e.g. better infanticide protection) or indirect (i.e. better genes for their offspring; [39]) fitness benefits of these male traits to females [40, 41].

## 2 Methods

### 2.1 Study system

We studied a natural population of mandrills established in 2002 by the release of 36 captive individuals followed by the release of another 29 individuals in 2006, in the Lékédi park, a private park located in Southern Gabon [42]. Starting in 2003, wild males joined the group to reproduce. In early 2012, the Mandrillus Project was set-up to study this population, benefiting from an initial habituation of these captive-born individuals to human presence. In early 2020, only 8 females from ca. 210 individuals were captive-born. All individuals were individually-recognized, daily monitored and censused.

### 2.2 Behavioural data

Trained observers, blind to the topic of this study, collected daily ad libitum behavioural observations and performed 5-min focal sampling on all study individuals [43]. In this study, we used 2182 hours of focal data collected on 81 adult females aged ≥4 yrs (mean±SD: 26.9±39.3h per female) and 670 hours collected on 34 subadult and adult males aged ≥9 yrs (19.7±29.2h per male), collected from August 2012 to March 2020. We included subadult males (aged 9-10 yrs) because they have usually reached their full adult body size [44] and have started competing with other males and mate with adult females [45]. During focal sampling, sexual and agonistic interactions between a focal individual and its groupmates were recorded. The observers systematically recorded copulations of males with females (n=275). Male aggressive events towards females included grasping/hitting (n=401), biting (n=18), chasing (n=65), lunging (n=383), slapping the ground (n=138) and head bobbing (n=567). For the analyses below, we ran the models including all these behaviours and we also replicated the analyses using only severe aggression (grasping/hitting, biting and chasing) or only threats (lunging, slapping the ground and head bobbing) because both categories produce different female behavioural reactions (see discussion). Dominance ranks were established separately for each sex (on a yearly basis for females and on a monthly basis for males) based on avoidance and displacements and calculated using normalized David’s score ([46]; as per [47]). Female rank is maternally inherited and generally stable during a female’s life [48]. Here, females were divided into three classes of equal size (high-, medium- and low-ranking) while male rank was considered as a binary variable (alpha versus non-alpha) because of the distinct behavioural characteristics of the alpha male, who monopolizes most swollen females and is relentlessly challenged by other males [49]. In the test for intimidation, in case the swollen period spanned over two consecutive months, a male was considered as alpha if he achieved the highest position for at least one of these two months.

### 2.3 Age and male immigration patterns

The exact date of birth was known for 25 individuals. For the remaining 90 individuals, the date of birth was estimated using body size, condition and patterns of tooth eruption and wear [50]. The error made when estimating the age of these 90 individuals was less than a year (50 individuals), two years (26 individuals), three years (13 individuals) or five years (1 individual). Long-term life-history and demographic data were also available from all individuals.

Census data allowed to reconstitute patterns of male residency in the group. Here, we considered a male as resident in a given mating season when censused in the group late during the preceding birth season, between January and March. When censused for the first time during the mating season (which takes place once per year between April and September) we considered the male as immigrant. For immigrant males, the first census date was the “arrival date”. Each year, the day of arrival of the first immigrant male in the group was considered as the onset of the mating season (figure S1).

### 2.4 Female reproductive state and sex ratio

During each female estrous cycle, the perineal swelling inflates for several days until reaching a maximal swelling size around ovulation. Swelling size remains maximal for a few days before deflating within a few days. We used a scale from 0 to 3 (by increments of 0.5) to evaluate the swelling size of each female on a near-daily basis. The reproductive state of each adult female was also recorded on a near-daily basis. Each female was classified as: “non-swollen” (i.e. non-fertile phase of the cycle that does not fall within the following three categories), “swollen” (i.e. with a perineal sexual swelling), “pregnant” (i.e. with a characteristic pregnancy swelling and/or if she gave birth 163-190 days afterwards (average gestation length: mean±SD: 175.0±4.7 days; [32]) or “lactating” (i.e. nursing a ≤6 month-old infant without having resumed cycling). Finally, females were considered as nulliparous until their first parturition, and parous afterwards. We calculated monthly adult group sex ratio (SR) or group operational sex ratio (OSR) as the number of females (for SR) or females with inflating sexual swelling or swelling of maximal size (for OSR) divided by the number of males aged 9 yrs and above that were censused in the group that month.

### 2.5 Injuries

We recorded the occurrence, type of wound, freshness and body location of any injury on a near-daily basis on all subjects [51]. A total of 90 injuries (limping n=15, puncture of the skin n=11, bleeding or swollen skin n=48, other n=16) were recorded on 43 females over the study period. For most injuries, we did not witness the interaction and the cause but in the three cases with a known context the injury was inflicted by a male. We never observed violent female-female aggression resulting in an injury.

### 2.6 Statistical Analyses

To test whether male aggression targets swollen females preferentially (first prediction), we ran a binomial generalized linear mixed models (GLMMs) with a logit link function to study the relationship between the probability that a female received aggression by any (adult or subadult) male during that female focal observation (0/1; response variable) and her reproductive state at the time of observation (non-swollen, swollen, pregnant and lactating; for sample sizes, see table S1). We further controlled for the following fixed effects: female dominance rank (high-, medium- or low-ranking) to test if higher-ranking females are preferentially targeted by males, parity (nulliparous or parous) to test if parous females are preferentially targeted by males, SR to test if the number of males in relation to females in the group influences the probability of occurrence of male aggression and the duration of focal observation (≤5min) to control for the observation time. Female identity and the year of focal observation were fitted as random factors. Second, we ran a similar model (same structure of fixed and random effects) with the response variable corresponding to the probability that a female received aggression by groupmates other than adult or subadult males. By doing so, we tested if swollen females were generally more targeted than any other female, regardless of the age-sex group of the aggressor.

To test whether swollen females were more injured than females in other states (second prediction), we ran a binomial GLMM with a logit link function to study the relationship between the probability that a female got injured (observed injured for first time) on a given day (0/1; response variable) and her reproductive state that same day. As above, we further controlled for the following variables: female dominance rank and parity, and SR. Female identity and the year of focal observation were fitted as random factors (table S1). The daily monitoring of the group allowed us to detect with accuracy the day of occurrence of each injury.

We then tested whether males who were more aggressive also had a higher mating probability with their victim (third prediction). To study intimidation, we performed a binomial GLMM with a logit link function to test whether the rate of aggression received by a female from a given male (continuous fixed effect) before the next estrous cycle of the female increased the probability of copulation of that heterosexual dyad during the female’s swollen period (0/1; response variable). The “aggression window” before the swollen period was defined as the time elapsed between the onset of the mating season (for resident males) or a male’s arrival in the group a given year (for immigrant males) and until the beginning of the swollen period of the female (spanning from the first day of a female’s sexual swelling to the last day where swelling size was maximal: mean±SD: 10.6±5.1 days; figure S1). We pooled focal observations from females and males (table S1). We controlled for the following fixed effects in our model: female dominance rank and parity, OSR (since we focused only on swollen females for that prediction) in the month corresponding to the first day of maximal swelling, male dominance rank (alpha vs. non-alpha) that same month in interaction with the rate of male aggression (to test whether the aggression of alpha males had a greater impact on their mating success than the aggression of subordinate males) and the total focal observation time of the studied heterosexual dyad (during the swollen period of the female) to control for the time of observation. Female identity, male identity and year of observation were fitted as random factors. We restricted our analyses to those heterosexual dyads that were observed for at least 30 minutes of focal time during the female swollen period to avoid biases due to under-sampling that would prevent us from estimating reliably mating probability. However. we validated that our results remained similar when we used slightly different thresholds (25 or 35 minutes) or no threshold at all. We further ran the same model but restricting the swollen period to the few days of the cycle during which the female was maximally swollen (i.e. where the probability of conception is the highest; mean±SD: 2.9±2.9 days). Finally, to test for immediate effects of male aggression, we ran the same model as above considering the rate of aggression received by a female from a given male during her swollen period only (figure S1, top line).

To test for sexual harassment, we assessed for each female and male focal observation during which an aggressive event was recorded from a male to a swollen female, whether a copulation occurred or not between that same heterosexual dyad in the 150 seconds following the aggression (see electronic supplementary material; figure S2). To test for male punishment, we assessed for each female and male focal observation during which a copulation event was recorded between a male and a swollen female, whether an aggression from a different male occurred towards the copulating female in the 150 following seconds (figure S2; table S1).

We further ran GLMM with a negative binomial distribution to test whether alpha males were more aggressive than subordinates during the mating season. We used as a response variable the number of aggression events a male directed towards all adult females during each month of the mating season (April to September). We considered only aggression towards females that were potential mating partners for males: late lactating females (during the 5th and 6th month of lactation when some females have already resumed cycling; MJEC personal observation), “non-swollen”, “swollen” and early pregnant females (during the first two months of pregnancy, since males may not be able to distinguish early pregnant from “non-swollen” females). We pooled focal observations from females and each given male (table S1). We included the following explanatory variables: male dominance rank (alpha vs. non-alpha) and age (to test if younger males are more aggressive) and the OSR (to test if males are more aggressive when there are few swollen females in comparison to the number of males in the group). The observation time of a given male and all the females was log-transformed, and fitted as an offset variable. Male identity and the year of observation were fitted as random factors.

We explored an alternative scenario to sexual coercion, the “aggressive male phenotype” hypothesis [39, 52], to test whether males with aggressive phenotypes have higher mating success than less aggressive males, potentially because aggression may act as a sexually selected trait and may be chosen by females. We reran the GLMM used for testing the occurrence of intimidation, including as an explanatory variable the overall rate of aggression directed by the focal male towards any groupmate (except for adult females) during the corresponding mating season.

We ran all the above statistical tests in R version 4.0.3. For generalized linear mixed models (GLMMs; summarized in table S1) we used the glmer function of the lme4 package [53] (binomial models) and glmmTMB from the package glmmTMB [54] (negative binomial model). Whenever a singular fit was observed, we reran the relevant model with the bglmer function of the blme package [55]. Whenever necessary we increased the number of iterations and/or we changed the optimizer of the model to achieve model convergence of the model and improve its fit. We used the Anova function of the car package [56] to test for the significance of fixed factors and computed their 95% confidence intervals. We further used the vif function of the same package to detect multicollinearities. All VIFs were <2.5 indicating no serious multicollinearities [57]. For multilevel categorical factors such as reproductive state, we switched the reference category sequentially [58] in order to test for pairwise differences between categories. We explored the distribution of residuals to validate the models using the functions testDispersion and simulateResiduals from the DHARMa package [59].

## 3 Results

### 3.1 Prediction 1: Male aggression targets swollen females

Swollen females received significantly more aggression from males (mean±SD: 0.613±1.070 bouts per hour) than females in any other reproductive state (non-swollen: 0.331±0.661, pregnant: 0.309±0.528 and lactating: 0.288±0.562; figure 1a, table 1). Such pattern was found for both severe aggression (rate toward swollen females: 0.349±0.948 bouts/hour, Chisq=12.539, p-value=0.006) and threats (0.260±0.390 bouts/hour, Chisq=8.660, p-value=0.034). By contrast, swollen females were not significantly more targeted by other groupmates (figure S3, table S2). In addition, high-ranking females received more male aggression than lower-ranking females (high-ranking females: 0.461±0.328 bouts/hour, medium-ranking females: 0.216±0.240, low-ranking females: 0.148±0.149, table 1).

**Table 1:** Male aggression in relation to female reproductive state (for sample sizes, see table S1). Significant p-values and confidence intervals that did not cross zero appear in bold. The significance of each variable was assessed using chi-square tests (Chisq), while the significance of each level of a categorical variable was evaluated against a reference level (noted ‘Ref’) according to whether their confidence intervals (CI) overlap or not.

| Response variable: Probability of receiving aggression from adult males (0/1) |  |  |  |  |  |
| --- | --- | --- | --- | --- | --- |
| Fixed Factor | Level | Estimate | CI 95% | Chisq | P-value |
| Reproductive State | Swollen (Ref: Non-Swollen) | 0.442 | **[0.170;0.714]** | 15.926 | **0.001** |
|  | Pregnant (Ref: Non-Swollen) | 0.070 | [-0.132;0.273] |  |  |
|  | Lactating (Ref: Non-Swollen) | -0.094 | [-0.309;0.122] |  |  |
|  | Swollen (Ref: Lactating) | 0.536 | **[0.268;0.804]** |  |  |
|  | Pregnant (Ref: Lactating) | 0.164 | [-0.030;0.358] |  |  |
|  | Swollen (Ref: Pregnant) | 0.372 | **[0.116;0.628]** |  |  |
| Female Rank | Low Rank (Ref: High Rank) | -0.718 | **[-0.981;-0.456]** | 31.124 | **<0.001** |
|  | Medium Rank (Ref: High Rank) | -0.554 | **[-0.904;-0.203]** |  |  |
| Female Parity | Parous (Ref: Nulliparous) | 0.150 | [-0.230;0.529] | 0.599 | 0.439 |
| Group Sex Ratio |  | -0.014 | [-0.059;0.031] | 0.375 | 0.54 |
| Observation Time |  | -0.097 | **[-0.167;-0.027]** | 7.459 | **0.006** |

**Figure 1:**
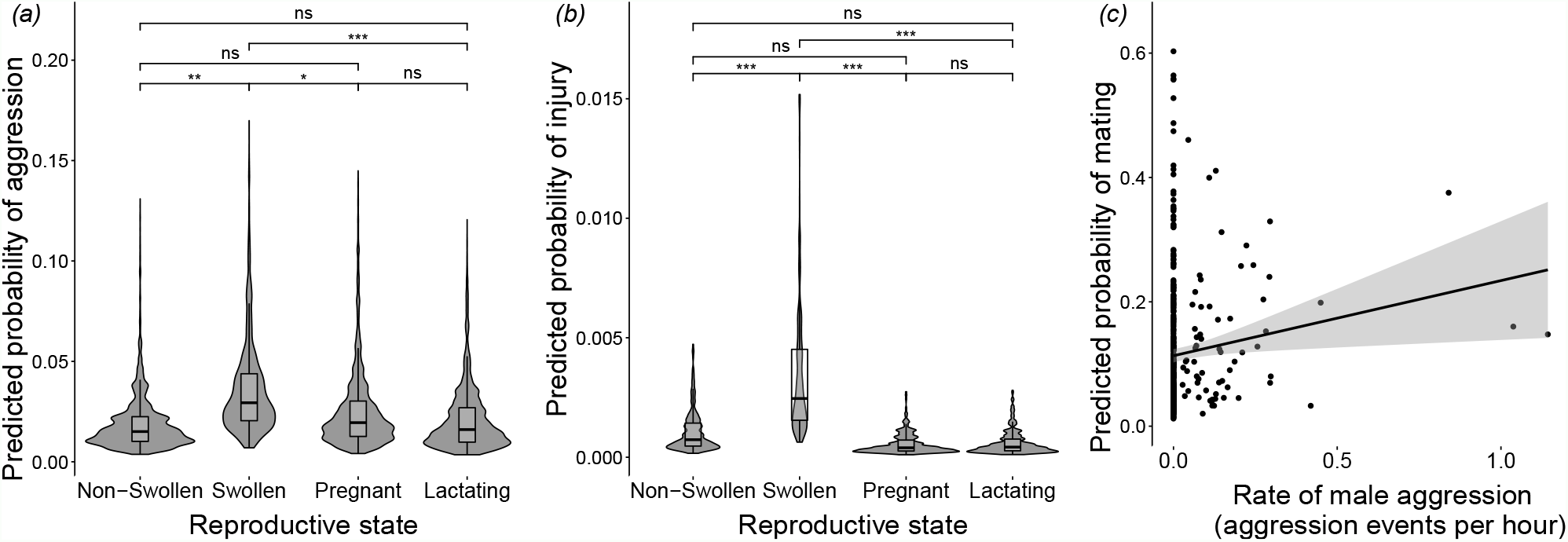
Results of the tests of the three predictions of the sexual coercion hypothesis. (a) Predicted probability of male aggression received by females as a function of their reproductive state. (b) Predicted probability for females to get injured as a function of their reproductive state. (c) Predicted probability of copulation of a heterosexual dyad as a function of male aggression rate (number of events per hour) received by the female before her swollen period. The fitted values of the GLMMs are shown on the y-axes. In a and b, the violin plots show the predicted probabilities while pairwise comparisons across female reproductive states with corresponding p-values are shown. ‘ns’: not significant (p>0.05); *: p<0.05; **: p< 0.01; ***: p<0.001. In c, the estimate and p-value are shown, while for graphical purposes, the regression line is simple linear fit and the shaded area shows the 95% confidence intervals.

### 3.2 Prediction 2: Swollen females are more injured

Swollen females were, on average, about five times more likely to become injured (mean±SD: 0.005±0.016 injuries per day) than females in any other reproductive state (non-swollen: 0.001±0.004, pregnant: 0.001±0.002 and lactating: 0.001±0.002; figure 1b). None of the other fixed factors, including female rank, parity and the group sex-ratio were significantly correlated with the probability of injury (table 2).

**Table 2:** Injuries in relation to female reproductive state (for sample sizes, see table S1). Significant p-values and confidence intervals that did not cross zero appear in bold. The significance of each variable was assessed using chi-square tests (Chisq), while the significance of each level of a categorical variable was evaluated against a reference level (noted ‘Ref’) according to whether their confidence intervals (CI) overlap or not.

| Response variable: Probability of having an injury (0/1) |  |  |  |  |  |
| --- | --- | --- | --- | --- | --- |
| Fixed Factor | Level | Estimate | CI 95% | Chisq | P-value |
| Reproductive State | Swollen (Ref: Non-Swollen) | 1.183 | **[0.579;1.787]** | 34.535 | <0.001 |
|  | Pregnant (Ref: Non-Swollen) | -0.452 | [-1.026;0.123] |  |  |
|  | Lactating (Ref: Non-Swollen) | -0.507 | [-1.076;0.061] |  |  |
|  | Swollen (Ref: Lactating) | 1.656 | **[1.013;2.299]** |  |  |
|  | Pregnant (Ref: Lactating) | 0.100 | [-0.503;0.704] |  |  |
|  | Swollen (Ref: Pregnant) | 1.556 | **[0.943;2.169]** |  |  |
| Female Rank | Low Rank (Ref: High Rank) | 0.203 | [-0.396;0.802] | 2.812 | 0.245 |
|  | Medium Rank (Ref: High Rank) | -0.418 | [-1.146;0.310] |  |  |
| Female Parity | Parous (Ref: Nulliparous) | 0.132 | [-0.826;1.090] | 0.073 | 0.787 |
| Group Sex Ratio |  | -0.013 | [-0.109;0.083] | 0.071 | 0.789 |

### 3.3 Prediction 3: Aggressive males have higher mating success with their victim

We found support for sexual intimidation in mandrills: the rate of male aggression received by a female during the time window preceding her swollen period (starting at the onset of a given mating season for resident males, or at male’s arrival date in the group for immigrant males) was significantly and positively correlated to the probability of copulation of the dyad during that swollen period (figure 1c, table 3). In dyads with no male aggression, the average number of copulation per observation time was 0.09±0.24 (±SD). By comparison, dyads where the male assaulted the female e.g. at least 0.1 times per hour, the average number of copulation per observation time doubled (0.17±0.45). Alpha males copulated more than subordinate males, while female rank, parity, OSR and the interaction between male rank and aggression (Chisq=0.030, p-value=0.862) were not significantly correlated with the probability of copulation (table 3). The correlation between male aggression and mating within dyads remained significant when restricting the swollen period to the few days where a female was maximally swollen (i.e. close to ovulation, Chisq=4.574, p-value=0.032). However, the rate of male aggression calculated during the swollen period of the female (instead of before) did not significantly predict the probability of copulation during that same swollen period (table S3a). This indicates that immediate aggression (i.e. during the swollen period) did not clearly influence female mating pattern, while previous aggressive interactions over a longer period (i.e. before the swollen period) did. The pattern of correlation between aggression and subsequent mating holds when only including severe aggression (table S3b) and becomes marginally non-significant when only including threats (table S3c). Note that the rate of severe aggression and the rate of threats a female receives from a male were moderately correlated (Kendall’s tau=0.28, p-value<0.001).

**Table 3:** Male aggression and mating success (for sample sizes, see table S1). Probability of copulation of a heterosexual dyad during a female’s swollen period in relation to the rate of male aggression received before that swollen period. Significant p-values and confidence intervals that did not cross zero appear in bold. The significance of each variable was assessed using chi-square tests (Chisq), while the significance of each level of a categorical variable was evaluated against a reference level (noted ‘Ref’) according to whether their confidence intervals (CI) overlap or not.

| Response variable: Mating during the swollen period (0/1) |  |  |  |  |  |
| --- | --- | --- | --- | --- | --- |
| Fixed Factor | Level | Estimate | CI 95% | Chisq | P-value |
| Aggression Rate |  | 1.591 | **[0.115;3.067]** | 4.466 | **0.035** |
| Male Rank | Alpha (Ref: Non-alpha) | 1.242 | **[0.490;1.994]** | 10.476 | **0.001** |
| Female Rank | Low Rank (Ref: High Rank) | 0.699 | [-0.186;1.584] | 2.664 | 0.264 |
|  | Medium Rank (Ref: High Rank) | 0.715 | [-0.645;2.075] |  |  |
| Female Parity | Parous (Ref: Nulliparous) | -0.454 | [-2.815;1.907] | 0.142 | 0.706 |
| Operational Sex Ratio |  | 0.024 | [-0.495;0.543] | 0.008 | 0.928 |
| Observation Time |  | 0.548 | **[0.221;0.875]** | 10.807 | **0.001** |

We did not find support for sexual harassment and punishment. Following aggression, females copulated immediately (i.e. within 150 seconds) with their aggressor in only three out of 38 total cases of aggression observed between a male and a swollen female. Similarly, males were never observed directing aggression to a female in the 150 seconds after she copulated with a rival male (out of 173 observed copulations). Those sample sizes precluded any further formal statistical testing of those hypotheses.

Alpha males were significantly more aggressive towards adult females. Indeed, an alpha male assaulted, on average, about 2 times more adult females (mean±SD: 0.05±0.07 bouts per hour) than a non-alpha male (0.03±0.06; figure S4; table 4). In addition, males were more aggressive (marginally significant effect; table 4) when there were more swollen females in the group in relation to males but male aggression did not depend on its age (table 4).

**Table 4:** Male rank and aggression (for sample sizes, see table S1). Male aggression towards adult females in the months of the mating season in relation to male rank, age and sex ratio. Significant p-values and confidence intervals that did not cross zero appear in bold. The significance of each variable was assessed using chi-square tests (Chisq), while the significance of each level of a categorical variable was evaluated against a reference level (noted ‘Ref’) according to whether their confidence intervals (CI) overlap or not.

| Response variable: Aggression during a month of the mating season |  |  |  |  |  |
| --- | --- | --- | --- | --- | --- |
| Fixed Factor | Level | Estimate | CI 95% | Chisq | P-value |
| Male Rank | Alpha (Ref: Non-alpha) | 0.610 | **[0.050;1.171]** | 4.552 | **0.033** |
| Male age |  | 0.050 | [-0.067;0.167] | 0.707 | 0.400 |
| Operational Sex Ratio |  | 0.315 | [-0.005;0.634] | 3.728 | 0.054 |

Lastly, we did not find evidence for a female preference for aggressive male phenotypes, as females were not more likely to mate with the most aggressive males of the group (see electronic supplementary material).

## 4 Discussion

We found support for all three core predictions of the sexual coercion hypothesis in mandrills. First, swollen females received significantly more male aggression than other females. Elevated aggression towards females around ovulation has been observed frequently in mammals, even in species where females dominate males socially (e.g. spotted hyena (*Crocuta crocuta*): [60]), suggesting that sexual coercion is widespread. Second, swollen female mandrills were significantly more injured than females in other reproductive states. Such injuries are most likely caused by males because aggression from other groupmates did not intensify during female sexual receptivity. Male aggression thus potentially causes important fitness costs in female mandrills, as shown in other mammals exhibiting sexual coercion (e.g. feral sheep (*Ovis aries*): [61]; bottlenose dolphins (*Tursiops cf. aduncus*): [62], chacma baboons: [14], chimpanzees: [63]). These fitness costs may push females to comply and copulate more with aggressive males to avoid conflict escalation and the associated risk of injury [64, 65]. Third, our analysis suggests that increased and repeated male aggression before the receptive period increases male mating success with the targeted female at times where she is most likely fertile. This correlation holds true both with severe aggression and non-physical threats, which are only moderately correlated. Most studies on sexual coercion have focused exclusively on severe aggression [14, 13] but our results indicate that male mandrills use a wide aggressive repertoire, including threats, to coerce females. In this species, male threats (such as head-bob or ground-slap) typically produce little immediate behavioural reactions in females, but could increase their sexual compliance with the aggressor when exerted repeatedly [28], especially when male-female power asymmetry is high, as in mandrills, which display one of the largest sexual dimorphism in primates.

The observed correlation between male aggression and mating success does not seem well-explained by alternative interpretations to sexual coercion, as we did not find evidence supporting a female preference for particularly aggressive males. Females could potentially use male aggression as a badge of status [13, 66] to infer male competitive abilities, which may provide females with direct or indirect benefits [40, 41]. However, in our data, variation in aggression rates among heterosexual dyads explain male mating success better than male general aggressiveness, suggesting that male mating success reflects relational properties more than male aggressive phenotype. It is further possible that male-female aggression rates directly reflect differences in male-female spatial proximity, where males would direct more aggression to females who would happen to stand around them. However, patterns of spatial ranging in social groups are far from random, and typically reflect the group social structure, in the form of differentiated relationships (e.g. spatial proximity is positively correlated to the strength of social bond in wild boars (*Sus scrofa*) [67]). In such context, male-female aggressive rates are more likely to reflect the existence of such differentiated social bonds between males and females than a scenario where a male would attack females who randomly happen to stand in their proximity. In line with this, recent studies in chimpanzees indicate that males preferably coerce their affiliated female partners [68], mirroring observations in humans where intimate partner violence is extensive [69].

Our analyses reveal important aspects of the ecology of sexual coercion in mandrill societies. While we did not find evidence for sexual harassment, our results suggest that repeated aggression over extended periods increases mating probability to aggressors once females become fertile, and may further encourage them to stay around males who mate-guard them, as observed in hamadryas baboons (*Papio hamadryas*; [28]). Sexual intimidation has previously been shown in chimpanzees and chacma baboons [13, 14], two species characterized by relatively high male violence towards females. We found that male mandrills use severe aggression towards swollen females more often on average than chacma baboons (mean±SD: 0.350±0.950 vs 0.130±0.190 times per hr; [14]) and at a rate that lies high within the chimpanzee’s reported range [13, 63]. Such frequent use of coercion by mandrill males may relate to the fact that - unlike chimpanzees and chacma baboons - they breed seasonally, thus have a limited time window to achieve mating. Yet, swollen female mandrills are injured ca. three times less on average than chacma baboons (mean±SD: 0.005±0.016 vs 0.014±0.022 injuries per day; [14]). Hence, although male to female aggression is more frequent in mandrills than in chacma baboons, violent aggression resulting in serious injuries is probably less common.

Moreover, the fact that we did not find any evidence of punishment, likely reflects the absence of exclusive mating bonds in mandrills (outside mate-guarding episodes) and the ability of females to sneakily escape male monopolization strategies in their dense habitat. Punishment by males in response to female sexual activity with a rival has, for instance, been reported in geladas (*Theropithecus gelada*) which live in more open habitat [17] and where one leader male can aggressively defend sexual access to females from his family unit [**snyder-mackler2012concessionsa**]. To sum-up, our results are generally consistent with expectations based on the socio-ecology of mandrills, who (i) are highly dimorphic thus where males pay low costs of intersexual aggression, (ii) breed seasonally, and where males face high pressure to mate in a relatively short period, and (iii) live in a polygynandrous mating system, and where males and females form differentiated social bonds - allowing intimidation to function - but no exclusive mating bonds, preventing the use of punishment by males.

Male dominance status appeared influential in their coercive tendencies. Alpha male mandrills were more aggressive towards females during the mating season, and they copulated significantly more with females than non-alpha males. Given the high reproductive skew in favour of alpha male mandrills [34, 35], this result suggests that sexual coercion is an effective male reproductive strategy, although more detailed analysis is necessary in order to confirm the relationship between male coercion and reproductive success. Dominant males in other primates similarly use long-term coercive strategies to constrain female promiscuity and impose closer proximity (e.g. hamadryas baboons [28]). However, in other species, such as orang-utans, subordinate males have been reported to be more coercive, and use forced copulations more often than dominant males [27]. The use of coercive strategies may be rendered more difficult for subordinate males in group-living species compared to solitary ones, such as orang-utans, if other group members, including the alpha male, occasionally step in to defend the victim.

Our analyses further highlight that all females are not equally targeted by males. High-ranking females specifically receive more male aggression than low-ranking females, which may reflect male mating preferences because dominant females show better reproductive performances than subordinates [48, 32]. Similarly, male hyenas mate preferentially with high-ranking females [70, 71] while male chimpanzees direct more aggression towards parous than nulliparous females [13] and prefer old females [72], who have a higher rank and reproductive success than younger ones [73]. This result indicates that the highest costs of coercion are born by the most attractive females, as found in chimpanzees [13].

An important question remains whether and how female mandrills may navigate such a coercive landscape while still possibly expressing some mate choice [33]. Chimpanzee studies have raised contrasting results, with sexual coercion in some populations [13, 63] versus female mate choice in other populations [74, 75]. It is possible that such conflicting results reflect differences across populations, or alternatively methodological differences between studies, where studies of mate choice often measure female choice through differential rates of approaches of males by females [74], while studies of sexual coercion correlate aggression and mating rates [13, 14]. The growing body of work on sexual coercion generally casts doubts on inferring mate choice from rates of approaches [4], as such approaches, as well as any affiliative interaction, could instead reflect female attempts to appease coercive males (i.e. [65]). Alternatively, it’s possible that sexual coercion can co-occur with female mate choice, as is the case in humans.

Our work underlines the existence of sexual coercion in mandrills while evidence for female choice remains scarce in this species [33]. It is therefore hard, at this stage, to evaluate the freedom left for females to express their own reproductive strategies. Nevertheless, several mechanisms may help females to mitigate the constraints set by male coercion. They may form alliances with other females to defend themselves [3, 76] or heterosexual bonds with males who protect them [77]. They may also appease male aggressors to limit the risk of escalation and injuries [28, 65], fight-back against aggressors, flee, hide or close their genitals [78, 79]. Female mandrills may use some of these strategies, as their behavioural repertoire includes avoiding male approaches, laying down when males attempt to copulate with them, refusing some mating attempts [33, 37], interrupting copulation by fleeing away, seeking support from subordinate males against dominant ones (MJEC personal observation) or even forming violent coalitions against high-ranking males ([38], NS personal observation). In addition, previous studies on primates have demonstrated that female reproductive synchrony and large group sizes limit female monopolization by males (across species: [80]; in mandrills: [35]) and increase the potential for females to express their strategies, including mate choice or promiscuity [81, 82]. Therefore, the extreme size of mandrill social groups along with female reproductive synchrony, may facilitate the expression of female reproductive strategies and reduce male coercion.

Here we report new evidence for sexual intimidation in a species where males, despite being much larger than females, are not conspicuously aggressive towards them (at least from a human observer perspective). The temporal uncoupling between male aggression and copulation explains why sexual intimidation may have long been overlooked, while it increasingly appears influential at shaping the social structure and mating system of polygynandrous mammals [20].

## Acknowledgements

The present work would not have been possible without the work of past and present field assistants of the Mandrillus Project and the logistical support of SODEPAL-COMILOG society. This is a Mandrillus Project publication number *n*^o^26 and ISEM 2022-132 SUD. Version 5 of this preprint has been peer-reviewed and recommended by Peer Community In Ecology (https://doi.org/10.24072/pci.ecology.100099).

## Ethics

All applicable international, national, and/or institutional guidelines for the care and use of animals were followed. This study was approved by the CENAREST institute (permit number, AR003/20/MESRSTT/CENAREST/CG/CST/CSAR) and adhered to the legal requirements of Gabon for the ethical treatment of non-human primates.

## Data accessibility

The datasets and scripts necessary to replicate analyses included in this article are deposited in the public depository: https://doi.org/10.5281/zenodo.6607803

## Supplementary information availability

The supplementary material relevant to this article, is deposited in the following public repository: https://doi.org/10.1101/2022.02.07.479393

## Authors’ contributions

N.S., M.J.E.C., and E.H. designed the study; B.R.T. and P.A.R collected behavioural data; N.S. performed the statistical analyses; N.S., M.J.E.C., E.H. wrote the original draft and all authors critically contributed to the draft and approved submission.

