## Supplementary materials for "Sexual coercion in a natural mandrill population"

### Methods

#### Sexual harassment and sexual punishment

To study sexual harassment and sexual punishment we used only 5 minutes-long (complete) focal observations performed on adult swollen females or adult males. To test for sexual harassment, we assessed whether a male directed aggression towards a swollen female during the first half of the focal observation (150 seconds). If aggression occurred ('post-aggression observation'; figure S2a, middle line), we assessed whether a copulation occurred between the female and the male within a 150 second-time window following this aggression. We planned to use a post-conflict matched-control statistical framework to test whether females were more likely to copulate with a male immediately after he attacked her versus in absence of aggression from him. In short, we had planned to match each post-aggression observation with matched-control observations (i.e., observations of the same individuals in which no male aggression occurred during the first half; figure S2a) and compare the likelihood of copulation in those two different contexts. The time span of 150 seconds was chosen as the maximum length allowing post-aggression and matched-control observation to be of equal length. Similarly, to test for sexual punishment we assessed whether a male copulated with a swollen female during the first half of the focal observation. If a copulation occurred ('post-copulation observation'; figure S2b, middle line), we assessed whether an aggression from another male towards the copulating female was observed within a 150 second-time window following the copulation. We had planned to use a similar post-copulation matched-control analysis to test whether females were more likely to be attacked in the post-copulation observations versus in the matched-control observations without copulation. However, since we found few or no instance(s) of post-conflict copulations and post-copulation aggression, we did not pursue those analyses and only report raw data.

#### Testing the "aggressive male phenotype" hypothesis

We explored an alternative scenario to sexual coercion, the "aggressive male phenotype" hypothesis, by testing whether males that are more aggressive towards any groupmate are also those that copulate the most because aggression may act as a sexual trait chosen by females. We reran the same GLMM as the one used for testing the occurrence of intimidation, including as an explanatory variable, in addition to the aggression towards the given female, the rate of the overall aggression the male directed towards all groupmates except adult females during the corresponding mating season. Such overall aggression was quantified as the number of aggression events initiated by a given male towards any non-adult female group member divided by the total time of observation of this male during a given mating season.

The overall aggression displayed by males towards non-adult females did not influence their copulation success with adult females suggesting that females do not copulate more with the most aggressive males of the group, but with males that have been the most aggressive to them before their fertile (swollen) period. In particular, in the model including both aggression rates (overall and dyadic), the aggression rate towards all groupmates except adult females was not significant (Chisq=2.12, p-value=0.15) but the rate of aggression towards the dyad female was marginally significant (Estimate=1.529, CI95%=[-0.039;3.097], Chisq=3.654, p-value=0.056) in comparison to the model without the overall aggression rate where the dyadic aggression rate was clearly significant (Table 3).

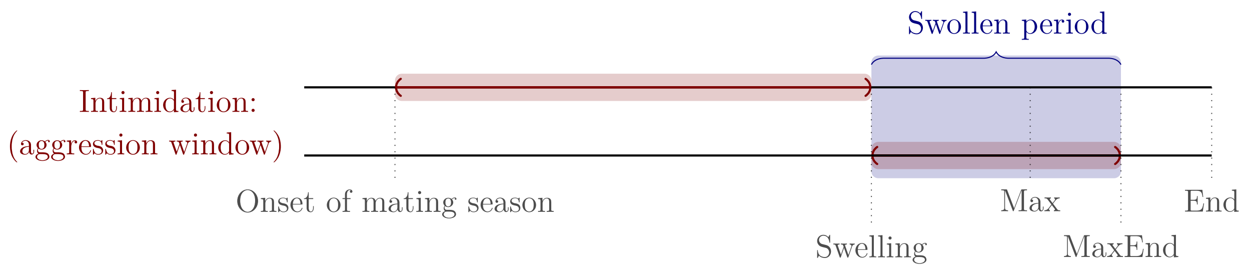


**Figure S1**: Analytical design for the tests of sexual intimidation. The swollen period is shown in blue, and the "aggression windows" are depicted in red. The top line represents the original test of intimidation. The bottom line represents the test with the alternative "aggression window". On the horizontal axis, the relevant temporally consecutive events (from left to right) are depicted (the distances among them can contextually fluctuate considerably). "Onset of mating season": onset of mating season (for residents) or arrival in the group (for non-residents), "Swelling": onset of the swollen period of the female, "Max": onset of the maximal swollen period of the female, "Max end": end of the swollen period (the last day of maximal swelling) and "End": complete deflation of the sexual swelling that started in "MaxEnd".

a

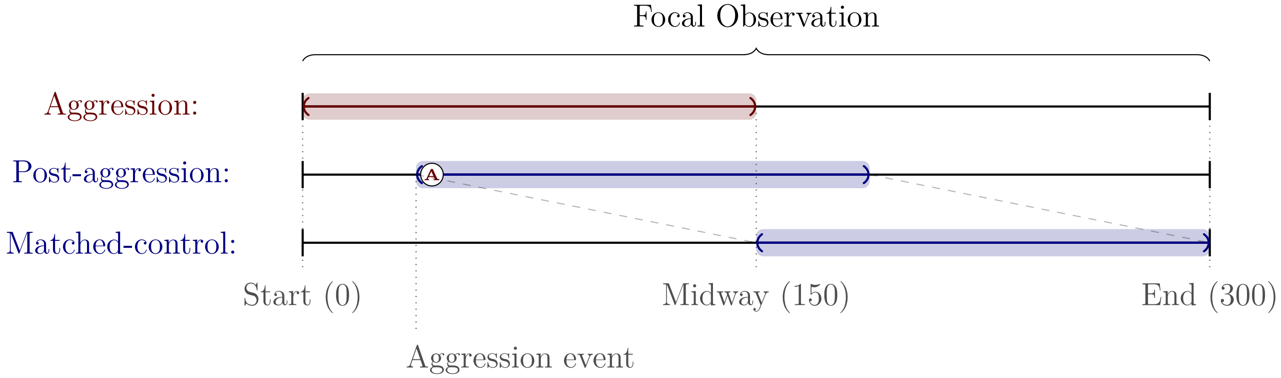


b

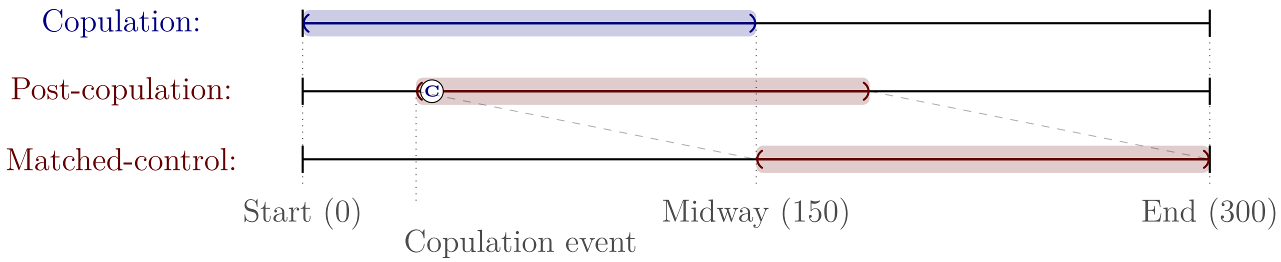


**Figure S2**: Analytical design for the planned test of sexual harassment and sexual punishment. (a) For each female or male focal observation, if an aggression occurred from a male towards a swollen female in the first 150 seconds (red), we assessed whether copulation of the same heterosexual dyad occurred within the 150 seconds following the aggression (post-aggression observation, blue – middle line); for the matched-control observations where no aggression occurred in this dyad in the first 150 seconds of the focal, we assessed whether a copulation was observed within the dyad during the last 150 seconds of the focal observation (blue – bottom line). (b) For each female or male focal observation, if a copulation occurred between a male and a swollen female in the first 150 seconds (blue), we examined whether aggression from another male towards the copulating female occurred within the 150 seconds following the copulation (post-copulation observation, red – middle line); for the matched-control observations where no copulation occurred in this dyad in the first 150 seconds of the focal, we assessed whether an aggression was observed during the last 150 seconds of the focal observation (red – bottom line).

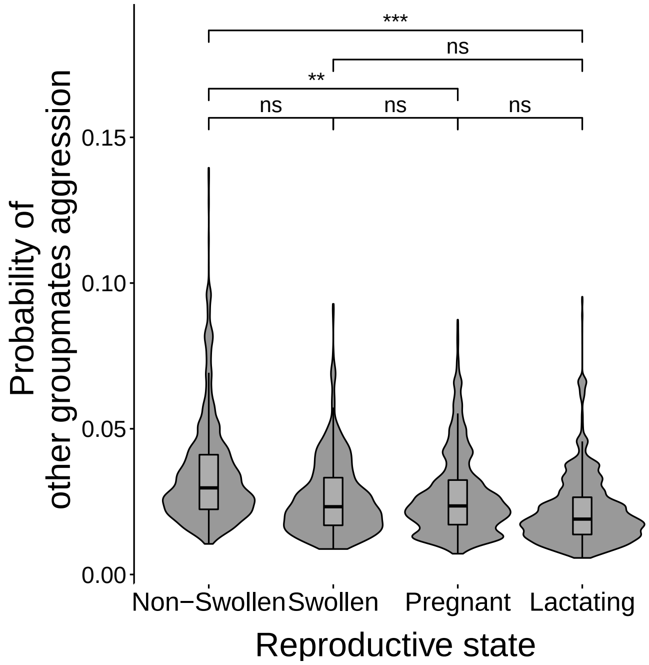


**Figure S3**: Predicted probability of aggression received by adult females from groupmates other than adult males in relation to female reproductive state. The fitted values of the GLMMs are shown on the Y-axis. The violin plots show the probability density. Pairwise comparisons across female reproductive states and corresponding p-values are shown. ‘ns’, not significant: p>0.05; *: p<0.05; **: p<0.01; ***: p<0.001.

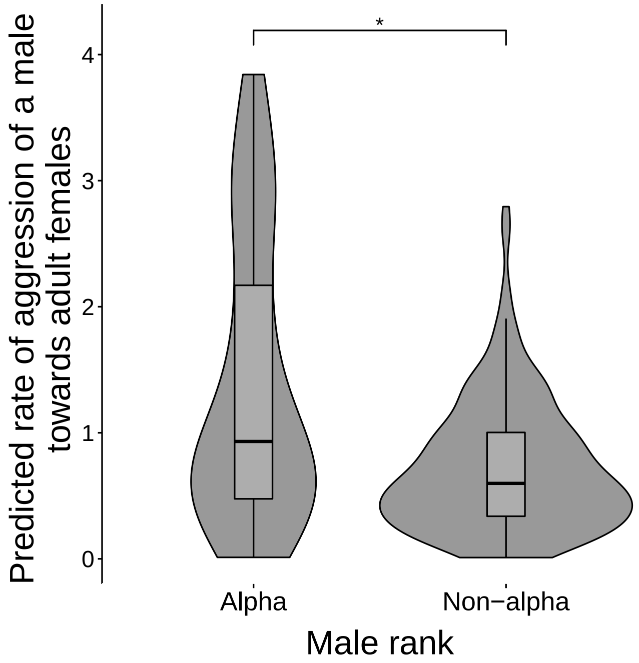


**Figure S4**: Predicted aggression rate of males towards adult females in relation to male rank (alpha vs non-alpha). The fitted values of the GLMM are shown on the Y-axis. The violin plots show the probability density. Pairwise comparisons across female reproductive states and corresponding p-values are shown. ‘ns’, not significant: p>0.05; *: p<0.05; **: p<0.01; ***: p<0.001.

**Table S1**: Summary of predictions and analyses with relevant sample size, figures and tables. All models followed a binomial distribution. S: swollen, NS: non-swollen, P: pregnant, L: lactating.

| Prediction | Resp. Variable | Sample | Fixed factors | Random Factors | Tabs & Figs |
|---|---|---|---|---|---|
| Swollen females receive more male aggression (1st prediction) | Probability of receiving male aggression during a focal observation | 31633 focals (2113h) on 80 females | Reproductive state (NS, S, P, L)<br>Female rank<br>Female parity<br>Group sex ratio | Female identity<br>Year of observation | Figure 1a<br>Table 1 |
| Swollen females do not receive more aggression from other groupmates | Probability of receiving other groupmate aggression during a focal observation | 31633 focals (2113h) on 80 females | Reproductive state (NS, S, P, L)<br>Female rank<br>Female parity<br>Group sex ratio | Female identity<br>Year of observation | Figure S4<br>Table S2 |
| Swollen females are at higher risk of injury (2nd prediction) | Probability of having an injury | 116.291 female.days (79 females and 2712 days) | Reproductive state (NS, S, P, L)<br>Female rank<br>Female parity<br>Group sex ratio | Female identity<br>Year of observation | Figure 1b<br>Table 2 |
| Male aggression increases male mating success (3rd prediction)<br>&<br>Aggressive phenotype hypothesis | Probability of copulation during the relevant period | Harassment/ Punishment: 1023 focals (85h) on 55 swollen females & 3590 focals (299h) on 34 males<br>Intimidation: 16212 focals (1116h) on 79 females & 5178 focals (366h) on 33 males | Male aggression (during the relevant period, towards the relevant individuals)<br>Female rank<br>Female parity<br>Operational sex ratio<br>Male rank (in interaction with aggression) | Female identity<br>Male identity<br>Year of observation | Figure 1c<br>Figure S1<br>Figure S2<br>Table 3<br>Table S3<br>Table S4 |
| Alpha males are more aggressive towards females | Aggression towards adult females | 16212 focals (1116h) on 79 females & 5178 focals (366h) on 33 males | Male rank<br>Male age<br>Operational sex ratio | Male identity<br>Year of observation | Figure S4<br>Table S4 |

**Table S2**: Aggression from other groupmates and female reproductive state. Significant p-values and confidence intervals (CI) that did not cross zero appear in bold. The significance of each variable was assessed using chi-square tests (Chisq), while the significance of each level of a categorical variable was evaluated against a reference level (noted 'Ref') according to whether their confidence intervals overlap or not.

| Response variable: Probability of receiving aggression from other groupmates (0/1) | | | | | |
|---|---|---|---|---|---|
| Fixed Factor | Level | Estimate | CI 95% | Chisq | P-value |
| Reproductive State | Swollen (Ref: Non-Swollen) | -0.192 | [-0.474;0.090] | 21.386 | **<0.001** |
| | Pregnant (Ref: Non-Swollen) | -0.241 | **[-0.412;-0.070]** | | |
| | Lactating (Ref: Non-Swollen) | -0.432 | **[-0.618;-0.246]** | | |
| | Swollen (Ref: Lactating) | 0.239 | [-0.056;0.534] | | |
| | Pregnant (Ref: Lactating) | 0.191 | [0.003;0.379] | | |
| | Swollen (Ref: Pregnant) | 0.049 | [-0.234;0.331] | | |
| Female Rank | Medium Rank (Ref: High Rank) | 0.256 | **[-0.090;0.602]** | 17.765 | **<0.001** |
| | Low Rank (Ref: High Rank) | 0.578 | [0.308;0.847] | | |
| Female Parity | Parous (Ref: Nulliparous) | -0.352 | **[-0.683;-0.021]** | 4.347 | **0.037** |
| Group Sex Ratio | | -0.024 | [-0.062;0.014] | 1.533 | 0.216 |
| Observation time | | 0.028 | [-0.041;0.098] | 0.647 | 0.421 |

**Table S3**: Male aggression and mating success (intimidation; alternative "aggression window"). (a) Probability of copulation of a male-female dyad during female's swollen period in relation to the rate of aggression received from the male during the female's swollen period. Probability of copulation of a male-female dyad during female's swollen period in relation to the rate of (b) severe aggression or (c) threats received from the male before the female's swollen period. Significant p-values and confidence intervals (CI) that did not cross zero appear in bold. The significance of each variable was assessed using chi-square tests (Chisq), while the significance of each level of a categorical variable was evaluated against a reference level (noted 'Ref') according to whether their confidence intervals overlap or not.

| Response variable: Mating during the swollen period (0/1) | | | | | | |
|---|---|---|---|---|---|---|
| Test | Fixed Factor | Level | Estimate | CI 95% | Chisq | P-value |
| a. Aggression in swollen period | Aggression Rate | | 0.173 | [-1.016;1.363] | 0.082 | 0.775 |
| | Male Rank | Alpha (Ref: Non-alpha) | 1.261 | **[0.542;1.979]** | 11.819 | **0.001** |
| | Female Rank | Low Rank (Ref: High Rank) | 0.617 | [-0.233;1.467] | 2.030 | 0.362 |
| | | Medium Rank (Ref: High Rank) | 0.288 | [-0.985;1.560] | | |
| | Female Parity | Parous (Ref: Nulliparous) | 0.304 | [-1.675;2.282] | 0.090 | 0.764 |
| | Operational Sex Ratio | | 0.112 | [-0.373;0.597] | 0.205 | 0.65 |
| | Observation Time | | 0.461 | **[0.160;0.761]** | 9.030 | **0.003** |
| b. Severe aggression only | Aggression Rate | | 6.307 | **[0.927;11.686]** | 5.280 | **0.022** |
| | Male Rank | Alpha (Ref: Non-alpha) | 1.291 | **[0.531;2.050]** | 11.086 | **0.001** |
| | Female Rank | Low Rank (Ref: High Rank) | 0.737 | [-0.153;1.627] | 2.879 | 0.237 |
| | | Medium Rank (Ref: High Rank) | 0.724 | [-0.639;2.087] | | |
| | Female Parity | Parous (Ref: Nulliparous) | -0.456 | [-2.843;1.931] | 0.140 | 0.708 |
| | Operational Sex Ratio | | 0.036 | [-0.494;0.565] | 0.017 | 0.895 |
| | Observation Time | | 0.537 | **[0.211;0.863]** | 10.446 | **0.001** |
| c. Threats only | Aggression Rate | | 2.111 | [-0.465;4.688] | 2.580 | 0.108 |
| | Male Rank | Alpha (Ref: Non-alpha) | 1.247 | **[0.511;1.983]** | 11.026 | **0.001** |
| | Female Rank | Low Rank (Ref: High Rank) | 0.682 | [-0.182;1.545] | 2.609 | 0.271 |
| | | Medium Rank (Ref: High Rank) | 0.673 | [-0.672;2.019] | | |
| | Female Parity | Parous (Ref: Nulliparous) | -0.510 | [-2.854;1.834] | 0.182 | 0.67 |
| | Operational Sex Ratio | | 0.005 | [-0.509;0.519] | 0.000 | 0.985 |
| | Observation Time | | 0.544 | **[0.220;0.868]** | 10.817 | **0.001** |
